# ProteoDUDes: Taxonomic profiling for metaproteomics with false positive reduction

**DOI:** 10.64898/2026.06.29.734936

**Authors:** Henning Schiebenhoefer, Thilo Muth, Stephan Fuchs, Bernhard Y. Renard

**Affiliations:** Hasso Plattner Institute, Digital Engineering Faculty, University of Potsdam, Prof.-Dr.-Helmert-Straße 2–3, 14482 Potsdam, Germany; Domain Specific Data Competence Centre (MF2), Department for Method Development, Research Infrastructure and Information Technology, Robert Koch Institute, 13353, Berlin, Germany; Genome Competence Centre (MF1), Department for Method Development, Research Infrastructure and Information Technology, Robert Koch Institute, 13353, Berlin, Germany; Software Architecture and Development Unit (IT4), Department for Method Development, Research Infrastructure and Information Technology, Robert Koch Institute, 13353, Berlin, Germany

## Abstract

Metaproteomics is the investigation of the protein composition of multi-organism samples. While metagenomics answers the question which organisms are present in a sample, metaproteomics additionally answers the question which organisms are active. State-of-the-art tools for annotating proteomic data with taxonomic information (e.g. Unipept, DIAMOND) do not control the false taxonomic identification rate, which can lead to incorrect results and thus incorrect interpretations, as we demonstrate with examples. ProteoDUDes processes the results from popular sequence annotation tools so that the proportion of true identifications in the result is at as high as or higher than in the the compared tools. We evaluate ProteoDUDes on simulated data and experimental mock community data. Our results indicate that ProteoDUDes has the same error rate as other tools on simulated data and half the error rate on the experimental mock community data. This allows more accurate statements to be made about which organisms are functionally active in a complex sample. ProteoDUDes is open-source and available at https://github.com/pirovc/dudes.

## Introduction

Taxonomic profiling is the identification and quantification of the microorganisms in a sample. Taxonomic profiling using metagenomics reveals the organisms present in a sample. It has been used, for instance, to characterize the present organisms in the human microbiome,^1^ microbiome differences between persons with higher and lower body weight,^2^ as well as characterizing ocean microbiomes across multiple locations across the globe. ^3^ On the other hand, metaproteomics reveals active organisms by investigating the proteins in a sample. Furthermore, metagenomics and metaproteomics have differing biases regarding enrichment protocols up front, data acquisition, and data analysis. These biases range from differing contamination risks during sample preparation^4^ to different organisms that are hard to identify (e.g. due to GC bias in sequencing^5^), to differing database biases.^6^

Metagenomics tools analyze marker genes or whole genome sequencing data. ^7,8^ Identifications in metaproteomics, on the other hand, are based on peptides, that is shorter sequences and a different alphabet with a larger spread of identified sequence lengths, different average lengths, or a larger dynamic range of expression levels that can span orders of magnitude.^9^ Therefore, transferring metagenomics tools to metaproteomics data requires additional effort to account for these differences.

An additional challenge faced in proteomics in general, but particularly in metaproteomics, is protein inference.^10^ A peptide, which is the actual unit of identification in proteomics, might be part of multiple proteins, thus hindering the inference of the originating protein. In metaproteomics, this problem is even more pronounced as related taxa exhibit similar protein sequences. This higher level of similarity of the genetic makeup complicates the distinction of taxa. This is further complicated by the fact that metaproteomics data is dominated by highly expressed proteins which tend to be better conserved.^9,11^

Metaproteomics often investigates environments that contain organisms beyond the well characterized present in reference databases and therefore uses e.g. translated metagenomes for peptide identification. Common approaches for the taxonomic annotation of proteomic data are the use of annotated protein databases (MPA^12^), peptide string matching (Unipept,^13^ MetaLab^14^), or protein sequence alignment (MetaGOmics,^15^ Prophane^16^). The use of annotated protein databases restricts the searchable protein sequences to known and annotated sequences. Peptide string matching searches identified peptide strings in a in-silico digested version of a reference database. Digestion rules for creating the database use trypsin as the digestion enzyme and only keep fully tryptic peptides, i.e. no missed cleavages which frequently occur in proteomics. Data analysis is therefore limited to tryptic digestion and fully tryptic peptides. For conserved proteins, string matching maps peptide sequences that originate from metagenomes to related organisms and thereby do low resolution (e.g. family instead of species) taxonomic annotation of uncharacterized species. Compared to the use of annotated protein databases, this allows the use of smaller, application specific protein sequence databases for proteomic searching. Sequence alignment searches identified sequences in reference databases and reports the best matching sequences as hits. In contrast to peptide string matching, sequence alignment takes all lengths of query sequences and thereby can align peptides with missed cleavages, regardless of the cleavage site. Commonly used tools for sequence alignment are DIAMOND^17^ and MMseqs2.^18^

In metagenomics, error control for taxonomic assignments is common practice.^19–21^ One tool for taxonomic profiling of metagenomic data is DUDes. ^22^ It controls the error of taxonomic annotations by executing overrepresentation tests for each candidate taxon in the data. Overrepresentation tests are executed iteratively from the root of the taxonomic tree towards the leaves, resulting in the deepest uncommon descendant (DUD). DUDes is limited to the analysis of alignment files in SAM format^1^ that contain the length of the reference sequence and cannot use UniProt reference databases, which are the default in metaproteomics.

A variety of tools have been proposed for metaproteomics, some ported from metagenomics to the challenges, some developed directly for metaproteomics (see^6^ for an overview). To the best of our knowledge, existing tools for taxonomic annotation of proteomic data rely on heuristic thresholds (e.g., e-value cutoffs) rather than explicit control of taxonomic annotation error. The taxonomic annotation depends largely on the utilized data analysis workflow.^6^ Hence, the reliability of taxonomic annotation is unknown making the interpretation of annotation results unreliable. There is therefore a need for a method that explicitly controls the error of taxonomic annotations.

We present ProteoDUDes, an extension of DUDes for taxonomic profiling of metaproteomic data against the UniProt database. ProteoDUDes can process data regardless of the cleaving enzyme and missed cleavages. To assess performance and limitations under controlled and experimental conditions, we validate the workflow on simulated and experimental mock community data of known taxonomic composition and show that it performs as well as or better than the competing tools Unipept and DIAMOND.

## Methods

We created ProteoDUDes by extending the DUDes^22^ tool to the inherent challenges of metaproteomic data processing. We enable taxonomic profiling of proteomic peptide data by adding support for UniProt^23^ reference databases and parsing of the output of common sequence alignment tools, see Figure 1. To avoid the protein inference problem, we chose to use peptides as the basic unit of identification. To evaluate taxonomic profiling, we analyzed benchmark datasets using ProteoDUDes, Unipept^13^ and DIAMOND.^17^ We picked Unipept and DIAMOND because they are easily automated and widely adopted based on their citation count. DIAMOND is used as a baseline for how ProteoDUDes compares against a best hit approach and Unipept as a benchmark. Two types of taxonomically defined benchmark data were used: simulated peptide datasets, constructed from UniProt protein sequences, and an experimental mock community dataset, derived from publicly available proteomic experiments. To assess performance, we compared each tools output with the respective samples ground truth at multiple taxonomic ranks.

**Figure 1:**
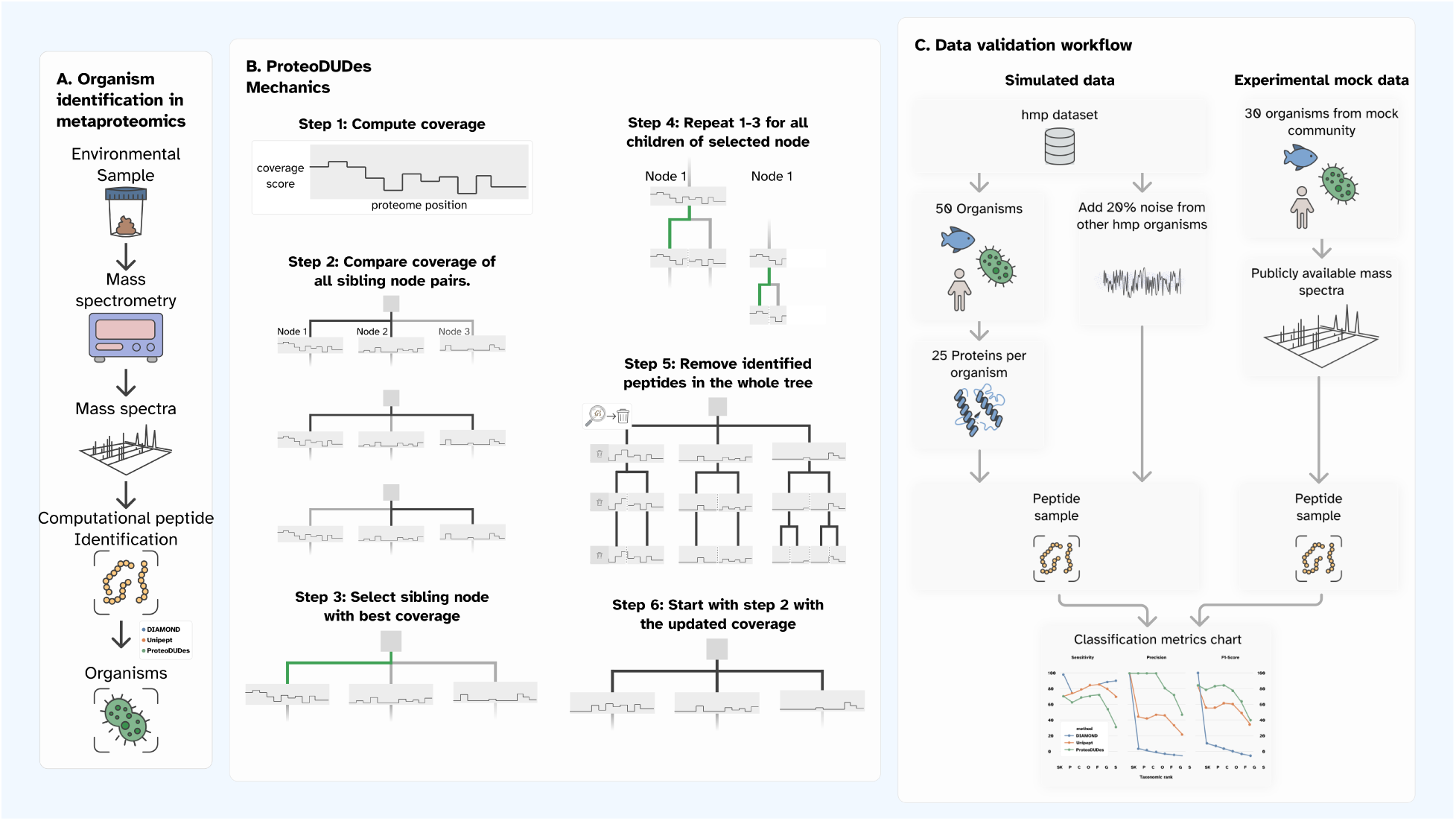
A. Organism identification in metaproteomics. An environmental sample, e.g. stool is processed and measured by a mass spectrometer. A proteomic search engine analyzes the generated mass spectra and assigns likely peptide sequences, optionally computing proteins that the peptides originated from. Finally, a taxonomic profiling tool annotates peptides or proteins with likely organisms of origin. *B. ProteoDUDes Mechanics* ProteoDUDes computes the proteome coverage for each taxonomic node that has any peptides mapped to it. By combining the coverage of all child nodes of each parent node, it computes the coverage on higher taxonomic ranks. In step 2, ProteoDUDes compares the coverage of all sibling nodes with each other and selects the best performing node for further analysis. This is repeated for the children of the selected node, until reaching a leaf of the taxonomic tree or a comparison yields no clear best performing node. ProteoDUDes reports the final thus identified deepest uncommon descendant (DUD) as identified and removes all peptides that belong to the identified node from all coverage plots. Since a peptide often occurs in multiple proteomes, this might affect not only the coverage of parent nodes of the identified node but also others. Finally, ProteoDUDes starts again at the top of the taxonomic tree and compares the coverage of all sibling nodes with each other, selecting the best performing node for further analysis. This process is repeated until convergence is reached. *C. Data validation workflow* We validate the ProteoDUDes workflow for taxonomic profiling in metaproteomics by analyzing data of known taxonomic composition. This data enables comparing the computed taxonomic profiles with the true taxonomic profiles and thereby assessing the perfomance. We generated simulated data by randomly drawing 50 organisms from the Human Microbiome Project (HMP) data set, drawing up to 25 proteins per organisms and finally drawing peptides with zero, one, and two missed cleavages from each protein. We complemented these peptides with noise peptides that we randomly drew from all accessions of HMP organisms. On top we used a experimental data set that consists of 30 organsisms. To generate these samples, a previous study mixed the organisms in a lab and subjected the generated sample to metaproteomic analysis. The generated mass spectra are publicly available and we use them for computational peptide identification and consecutive taxonomic profiling.

### ProteoDUDes

To perform taxonomic profiling on proteomic peptide data we created ProteoDUDes, an enhanced version of the DUDes^22^ tool (version 0.10.0). ProteoDUDes requires three inputs: (1) a set of query-to-reference-sequence matches, (2) a set of reference sequences, and (3) a taxonomic tree structure.

Usage of sequence alignment tools enables support for non-tryptic peptides, missed cleavages and sequence mismatches, all of which are not supported by string-based analysis tools. Popular tools for sequence alignment of proteomic data are DIAMOND^17^ and MMseqs2.^18^ To support the output of these tools, we added a custom query-to-reference-sequence format parser to ProteoDUDes. Additionally, we added support for reference matches in the popular UniProt database. The custom query-to-reference-sequence format is a tab separated file with the following columns: (1) query sequence identifier, identifying all rows that belong to the same peptide. (2) subject sequence identifier, identifying the UniProt reference sequence in the format “|{identifier}|”. (3) Subject sequence length, the length of the reference sequence. (4) Start of the alignment in the subject sequence. (5) E-value.

For the identification of taxons, ProteoDUDes assigns each query-to-reference-sequence match to the corresponding taxonomic node (see Figure 1, panel B). ProteoDUDes divides reference sequences of the entire reference database into equally sized bins and assigns each query-to-reference sequence match from the alignment tool output to the bin at the match position. ProteoDUDes uses the E-value from the alignment tool output to calculate a match score as follows: floor(−10 ∗ *log*_10_(E-value)). It sums up the scores of all matches in a bin to calculate the bin score, thus generating a coverage for each node (see Figure 1, panel B, step 1).

ProteoDUDes uses these coverages to compare nodes with each other, starting at the root of the taxonomic tree. It does a pairwise comparison of all nodes that share the same parent node (see Figure 1, panel B, step 2). For a single comparison, the tool calculates the cumulative bin score of each node and picks half of the bins of the node with the higher cumulative bins score. This number is chosen to normalize for differences in taxon size between nodes. It picks this number of highest scoring bins from both nodes. If the second node contains fewer bins than requested, the tool selects the number of bins available in the second node from both nodes. ProteoDUDes uses these bins to create a distribution of bin score differences between two taxons by repeatedly randomly assigning their bin scores to each taxon and calculating the difference in cumulative bin scores. If the actual bin score difference is larger than the specified significance threshold, the taxon with the larger score is deemed significant in this comparison. It repeats this process for all combinations of taxons on this taxonomic level that share a common parent. ProteoDUDes then counts the number of times a taxon is not significant and the number of times other taxons were significant against it. The taxon with the lowest total count is selected as identified. ProteoDUDes repeats this process for the direct child nodes of the identified taxon and compares them with each other (see 1, panel B, step 4). It repeats this process until there are no more taxonomic ranks left or no more identifications are possible. ProteoDUDes then filters out the identified query-to-reference-sequence matches (see 1, panel B, step 5) and continues with another iteration from the start of the taxonomic tree (see 1, panel B, step 6).

### Benchmarking

To assess the performance of taxonomic profiling tools, we require data of known taxonomic composition. By comparing the taxonomic profiles that the tools produce with the taxonomic profile of the input data, we can calculate the following performance metrics (see Figure 1, panel C).

#### Simulated Peptide Data

We simulated peptide data of known taxonomic composition, combined with a specified ratio of noise peptides. We chose peptides as the starting point to focus on false positives introduced by taxonomic annotation. We thus keep out errors introduced, by e.g. sample preparation, sample carry over in the instrumental analysis, or peptide identification. We chose the organisms from the expanded Human Microbiome Project (HMP) ^24^ as the basis of the simulation in order to make the taxonomic composition of our simulated samples reflect the taxonomic composition of the human microbiome.

Benchmark peptide samples of known taxonomic composition were created as follows: Species for generating simulated samples were taken from a previous study (, ^24^ supplementary table 2). 151 Species names were translated to taxonomic identifiers using the EMBL-EBI taxonomy API,^25^ resulting in 133 taxonomic identifiers. These were then mapped to protein accessions for 113 taxonomic identifiers using a UniProt idmapping file, version 2023 05. Protein sequences were retrieved from the UniProt API https://rest.uniprot.org/uniprotkb/.

For simulating signal peptides, 50 out of the 113 taxonomic identifiers were randomly sampled. For each taxonomic identifier, where available, up to 100 protein accessions were sampled. Protein sequences were subjected to tryptic digestion, with up to two cleavages. The number of peptides per protein was sampled from a poisson distribution with mean 10. Missed cleavages were weighted as follows: 0: 100, 1: 10, 2: 1.

False discoveries on the peptide and protein level in proteomics are most commonly controlled by applying a false discovery rate (FDR) cutoff by reverse database search, ^26^ which is an estimate for the ratio of false discoveries in the total number of discoveries. We argue that the effective rate of false discoveries is commonly higher than the stated FDR. An example supporting this claim is the fact that at a 1% FDR, which is an estimate for the ratio of identifications that is wrong, we would expect different tools to assign at most 2% different peptides to the same set of spectra. In reality, this non-overlap can be 15-20%.^27^ We therefore opted to add 20% noise peptides. We sampled these noise peptides from the accessions of all non-signal taxonomic identifiers in the HMP dataset. We restrict sampling of noise peptides to peptides not in the sample set.

Taxonomic lineages for the taxonomic ranks *superkingdom*, *phylum*, *class*, *order*, *family*, *genus*, and *species* were generated using the NCBI nodes.dmp file downloaded on 16th December 2023.

#### Experimental Mock Community Data

Experimental mock community data was taken from Kleiner et al.,^28^ PRIDE^29^ dataset PXD006118. This data is the largest biological sample with known taxonomic composition. Biological samples of organisms were mixed at specified ratios and subjected to proteomic analysis. We selected the raw data files with uneven protein amount (Run1_U1_2000ng.raw). The uneven mixture more closely represents the organism distribution in biological samples. The sample contains 30 organisms. The fasta file provided by Kleiner et al. was used as the proteomic search database (`Mock_Comm_RefDB_V3.fasta`).

Proteomic searching was executed using FragPipe (v21.1), with MSFragger 4.0,^30^ Ion-Quant 1.10.12,^31^ and Philosopher 5.1.0.^32^ Search parameters differing from the defaults are 10ppm precursor mass tolerance, a fragment mass tolerance of 0.1 Da and trypsin as digestion rule.

For further analysis, the FragPipe output file peptide.tsv file was used.

#### Evaluation

We calculate sensitivity to assess detection capability, precision to assess false positive control, and F_1_-score for assessing the trade-offs.

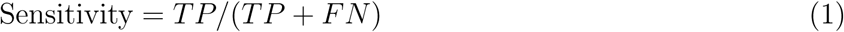

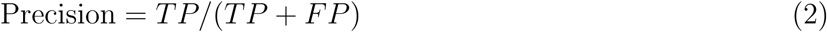

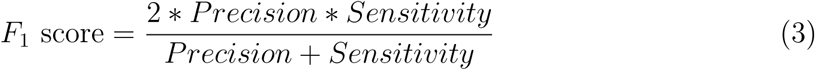

Where True Positives (TP) are the number of taxons that were identified as present and were in the sample, False Positives (FP) are the number of taxons that were identified but were not in the sample, and False Negatives (FN) are the number of taxons that were in the sample and were not identified as present. These metrics were calculated for each tool on each taxonomic rank of each analyzed sample.

#### Sequence Alignment

For taxonomic annotation, the peptide sequences were aligned with the UniProt database using DIAMOND^17^ (Version 2.0). UniProt Swiss-Prot and TrEMBL databases (Version 2023 05) were converted to a DIAMOND database and DIAMOND blastp was executed with the following parameters: `--fast --outfmt 6 qseqid sseqid slen sstart evalue`.

#### Unipept

The Unipept command line interface^33^ (Version 3.1.0) was used to annotate simulated and experimental mock community peptide sequences. The command line interface was executed using the following command: `cat {input} | prot2pept | peptfilter | unipept pept2lca -a -e`.

#### ProteoDUDes

ProteoDUDes was executed with the following parameters: The ProteoDUDes database was created using the dudesdb command in uniprot reference mode (-m up) on UniProt Swis-sProt and TrEMBL version 2023 05 containing 2.5 ∗ 10^8^ sequences and NCBI taxdump file downloaded on 16th December 2023. Identification was executed using the dudes command, providing input in custom blast file (-c) format. ProteoDUDes was executed on a machine with 2x AMD EPYC 7742, 64-Core processors and 1 TB RAM.

## Results and Discussion

We present a taxonomic profiling workflow for metaproteomics data. We evaluate the workflow on simulated data and a lab assembled mock community. We compare the results and computational requirements of our workflow with DIAMOND and Unipept. While ProteoDUDes is less sensitive than the compared tools, it outperforms them in precision, resulting in a higher F_1_-score, particularly on the arguably more relevant and realistic experimental mock community data.

### Computational resources

Sequence alignment and taxonomic profiling are resource heavy tasks, e.g. due to the large reference databases. Table 1 summarizes the peak memory usage and runtime of the database creation and sample analysis for each tool. Database creation resources vary widely between tools, with ProteoDUDes requiring nearly 200 times as much memory and 24 times as much runtime as DIAMOND does. Since we used the Unipept web service which uses a precomputed database, we do not include computational resources for the Unipept database creation. Each tool creates this database once, and reuses it for subsequent sample analyses. ProteoDUDes sample analysis requires 22 times as much memory and double the amount of runtime as DIAMOND for the largest sample. Unipept analyzes the samples on their remote hardware and therefore merely requires a web browser or the Unipept command line interface. ProteoDUDes requires specialized hardware such as a high performance computing environment or a workstation and analyzes a single sample in under 10 minutes.

**Table 1:** Computational requirements of each tool.

| Analysis Step | Tool | RAM [GB] | Time [hh:mm] |
| --- | --- | --- | --- |
| Database<br>Creation | DIAMOND | 5 | 00:33 |
|  | Unipept<sup>1</sup> | n/a | n/a |
|  | ProteoDUDes<sup>2</sup> | 945 | 13:00 |
| Sample<br>Analysis | DIAMOND<sup>3</sup> | 7 | 00:06 |
|  | Unipept<sup>4</sup> | minimal | minimal |
|  | ProteoDUDes<sup>3</sup> | 150 | 00:11 |

<sup>1</sup> Unipept database creation is carried out by Unipept operators.
<sup>2</sup> A precomputed database is available for download, see data availability section on page 17.
<sup>3</sup> maximum resource usage across simulated and experimental mock community data.
<sup>4</sup> Unipept sample analysis is run in a web browser.

ProteoDUDes has a large memory footprint, requiring 900 GB for database creation and 150 GB for sample analysis. The memory footprint during database creation is caused by ProteoDUDes loading the entire reference database into memory. During analysis, the largest contributor to memory usage is likely the data permutation. Hence, running ProteoDUDes requires a high performance computing environment or a workstation while DIAMOND can run on consumer computers, and Unipept can run on any device with internet access because it carries out the entire analysis at the Unipept servers. Unipept server based analysis restricts applicability due to data privacy concerns. A potential remedy for the large memory footprint during database creation is to process the database in chunks instead of as a whole. A precomputed database is available for download reducing hardware requirements https://zenodo.org/records/10680335/files/dudesdb_uniprot_202401.tar.gz.

Potential next steps are making ProteoDUDes less resource intensive during the database creation to make it future proof by still being able to process future, larger databases. By including reference lengths in the ProteoDUDes database, ProteoDUDes could process output from Unipept thereby allowing the integration into workflows that make use of Unipept.

### Simulated Peptide Data Analysis

While ProteoDUDes reuires large computational resources, any analysis can be locally performed. To evaluate the performance of ProteoDUDes against DIAMOND and Unipept, we first assessed the performance on simulated data (Figure 2). Simulated data consists of peptides sampled from 50 organisms from the HMP data, complemented with 20% noise peptides sampled from the remaining HMP organisms, representing false positive identifications. We measured performance using the metrics sensitivity, precision and F_1_-score across taxonomic ranks from *superkingdom* to *species*.

**Figure 2:**
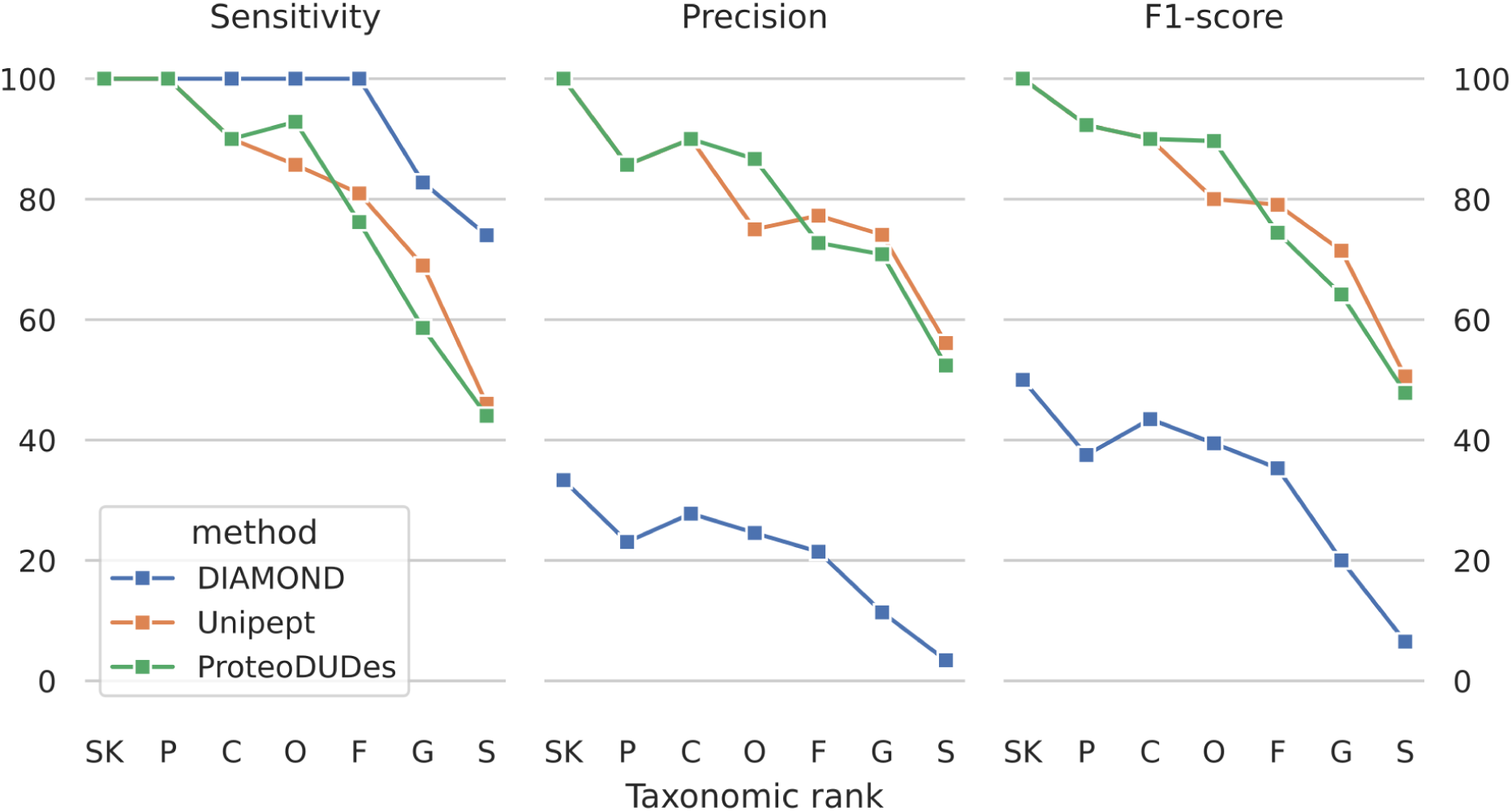
Performance of DIAMOND, Unipept, and ProteoDUDes on simulated Peptide Data. Data consists of 6,473 signal peptides sampled from 50 HMP organisms and 1,294 noise peptides, sampled from the non-signal HMP organisms. The subfigures show the metrics sensitivity, precision, and F_1_-score as percentage for each tool across the taxonomic ranks *SK: superkingdom*, *P: phylum*, *C: class*, *O: order*, *F: family*, *G: genus*, and *S: species*.

Sensitivity of tools ranges from 100% on *superkingdom* rank to 44% on *species rank*. Sensitivity is the highest for DIAMOND, with Unipept and ProteoDUDes being up to 30% less sensitive, particularly towards lower ranks such as *species*. In contrast to sensitivity, the spread of precision between the tools is much larger with ProteoDUDes and Unipept above 70% across all ranks except *species* and DIAMOND consistently at 33% or lower. Precision decreases for all tools towards lower ranks, with ProteoDUDes and Unipept decreasing to 52% and 56% on *species* rank, whereas DIAMOND drops to 3%. The small difference in sensitivity combined with the large differences in precision result in ProteoDUDes and Unipept having an at least 39% higher F_1_-score than DIAMOND across all ranks. Differences between ProteoDUDes and Unipept are below 10% across all metrics and ranks.

ProteoDUDes and Unipept have similar sensitivity, precision, and F_1_-score on simulated data. The simulated sequences originate entirely from UniProt and since Unipept uses UniProt as database, its string matching approach works well on the simulated data. Another potential reason is simulated data containing few missed cleavages, working to Unipepts advantage since it can only process fully tryptic peptides. In contrast, ProteoDUDes is not limited to fully tryptic peptides and can also analyze peptides with missed cleavages.

### Experimental Mock Community Data Analysis

We evaluated the performance of DIAMOND, Unipept, and ProteoDUDes on experimental mock community data (Figure 3). The data consists of 30 organisms, which were mixed at uneven mass ratios and subjected to proteomic analysis. This data is the largest experimental proteomic sample available with defined taxonomic composition. Similarly to the simulated dataset, performance metrics across taxonomic ranks from *superkingdom* to *species* are sensitivity, precision, and F_1_-score.

**Figure 3:**
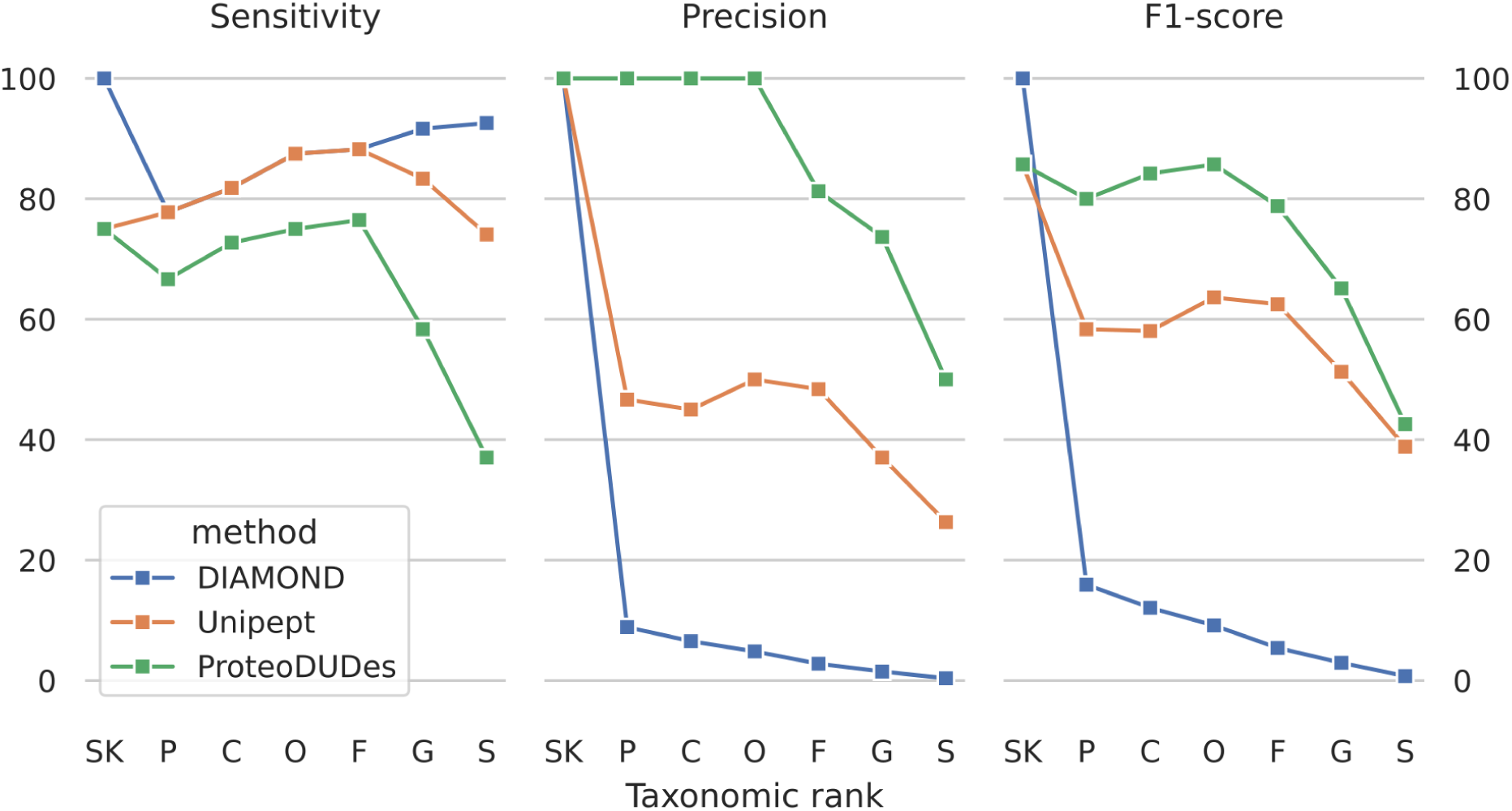
Performance of DIAMOND, Unipept, and ProteoDUDes on Experimental Mock Community Data. Mock Community Data consists 30 organisms which were mixed at uneven masses to approximate the protein composition of a biological sample. The subfigures show the metrics sensitivity, precision, and F_1_-score as percentage for each tool across the taxonomic ranks *SK: superkingdom*, *P: phylum*, *C: class*, *O: order*, *F: family*, *G: genus*, and *S: species*.

The sensitivity of the tools ranges from 100% to 37%. ProteoDUDes is up to 12.5% less sensitive than DIAMOND and Unipept down to *family* rank, thereafter the difference increases to 37% less than Unipept and 56% less than DIAMOND on *species* rank. However, ProteoDUDes has the highest precision of all evaluated tools, with 100% for the ranks *superkingdom* to *order*, and decreasing steadily to 50% on *species* rank from there. The large differences in precision combined with small differences in sensitivity between the tools lead to ProteoDUDes having the highest F_1_-score.

ProteoDUDes has a higher precision and F_1_-score than the competing tools on the experimental mock community data, which provide the closest approximation to real biological samples while retaining a defined taxonomic composition. While neither DIAMOND nor Unipept have any means of filtering out False Positives, ProteoDUDes can aggregate evidence across the taxonomic tree instead of relying on single best hits. Thus, ProteoDUDes can provide more precise taxonomic identifications than the other metaproteomics taxonomy profiling tools. The gain in precision is particularly high at intermediate ranks of the taxonomic tree and decreases toward the terminal nodes. Here, also ProteoDUDes struggles to always correctly distinguish between highly similar species given the extremely small number of spectra that are unique for one species in the analyzed datasets.

ProteoDUDes’ gain in precision over other tools decreases on more specific taxons and is gone on *species* rank. Since DIAMOND results provide the input of ProteoDUDes, a potential reason for the loss in precision is that DIAMOND reports all matches of each peptide, resulting in an increasing amount of False Positives when descending the taxonomic tree to more specific ranks. More False Positives lead to ProteoDUDes having to compare more and more nodes with each other when proceeding to more specific ranks, leading to lower statistical power and higher chances of identifying a node that has a high coverage by pure chance. When comparing Unipepts precision with DIAMONDs precision, we see the same trend. ProteoDUDes’ results on higher taxonomic ranks are more reliable than on lower taxonomic ranks. This effect might be alleviated by configuring the sequence alignment in a stricter way, sacrificing sensitivity for precision.

With metaproteomics moving beyond usage in fundamental research towards diagnostic applications,^34^ the need of precision in the taxonomic annotation becomes more pronounced. With its basis on permutation statistics and the usage of full coverage information, ProteoDUDes provides a more robust basis.

## Conclusions

We present a workflow for taxonomic profiling with increased precision compared to the state of the art tools. It increases precision by removing spurious hits with a transparent method, making taxonomic profiling of metaproteomic data more reliable. It supports any kind of proteolytic enzyme and can make full use of non-tryptic peptides. The integrated data simulation workflow allows validation of future taxonomic profiling methods.

## Acknowledgement

This work is supported by a European Research Council (ERC) grant (eXplAInProt, 101124385) to B.Y.R..

The authors declare no competing financial interest.

Author Contributions: H.S., T.M., S.F., and B.Y.R. conceived the method, H.S. and B.Y.R. conceived the validation strategy, H.S. conducted the validation and analyzed the results. H.S., T.M., S.F., and B.Y.R. wrote and reviewed the manuscript.

The authors thank Luciana Serna Wills assistance with illustrations, Vitor Piro for technical guidance, and Christoph Schlaffner for reviewing the manuscript.

## Data Availability Statement

ProteoDUDes is open-source and available at https://github.com/pirovc/dudes. A precomputed database is available at https://zenodo.org/records/10680335/files/dudesdb_uniprot_202401.tar.gz. The data validation workflow, starting with FragPipe output and comprising data simulation, sequence alignment, taxonomic profiling, and generation of evaluation plots, is available at https://github.com/rababerladuseladim/proteodudes-evaluation.

## TOC Graphic

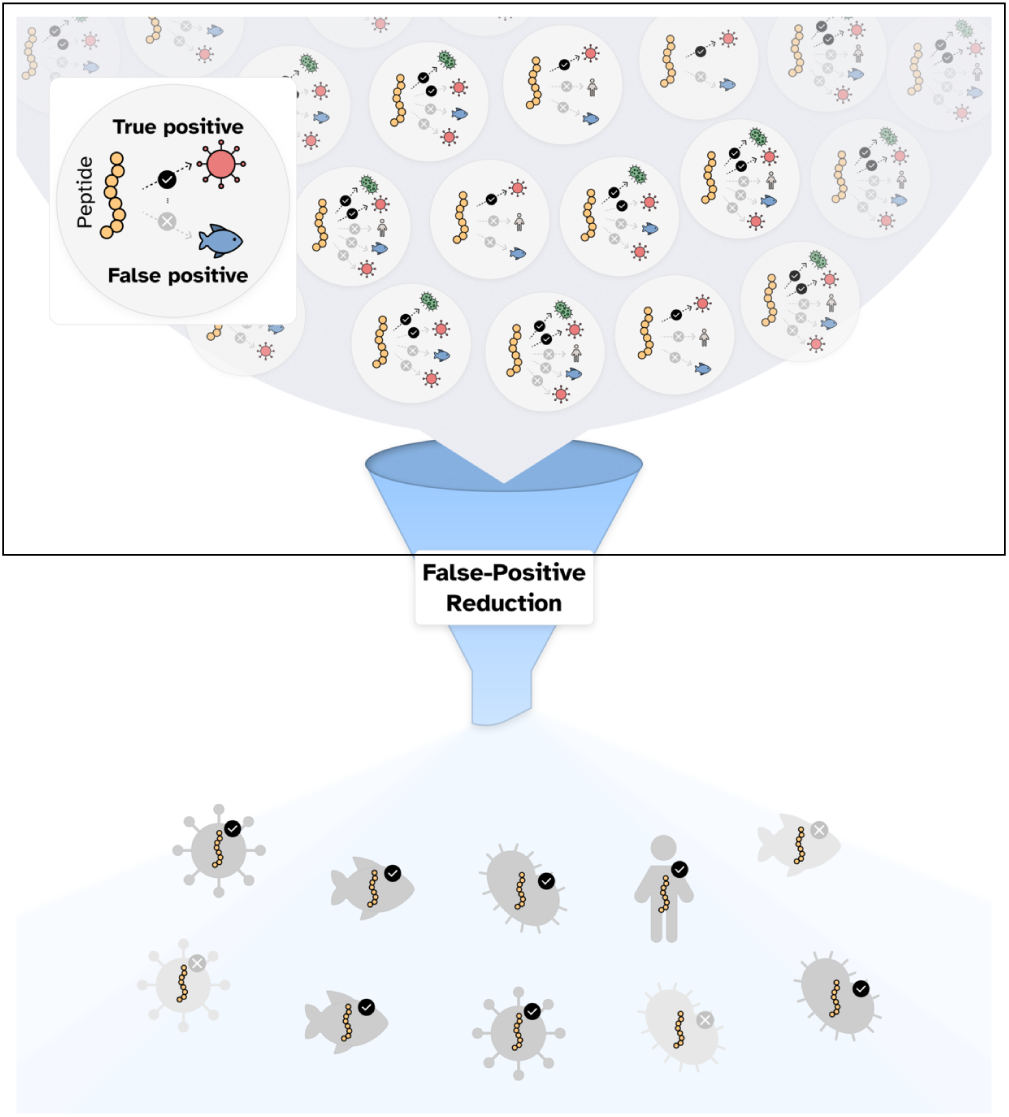

## Footnotes

1 https://samtools.github.io/hts-specs/SAMv1.pdf

## References

(1) Human Microbiome Project Consortium Structure, function and diversity of the healthy human microbiome. Nature 2012, 486, 207–214.

(2) Turnbaugh, P. J.; Ley, R. E.; Mahowald, M. A.; Magrini, V.; Mardis, E. R.; Gordon, J. I. An obesity-associated gut microbiome with increased capacity for energy harvest. Nature 2006, 444, 1027–1031.

(3) Sunagawa, S. et al. Structure and function of the global ocean microbiome. Science 2015, 348, 1261359.

(4) Piro, V. C.; Renard, B. Y. Contamination detection and microbiome exploration with GRIMER. GigaScience 2023, 12, giad017.

(5) Holcik, L.; Von Haeseler, A.; Pflug, F. G. Genomic GC bias correction improves species abundance estimation from metagenomic data. Nature Communications 2025, 16, 10523.

(6) Van Den Bossche, T., et al. Critical Assessment of MetaProteome Investigation (CAMPI): a multi-laboratory comparison of established workflows. Nature Communications 2021, 12, 7305.

(7) Wood, D. E.; Lu, J.; Langmead, B. Improved metagenomic analysis with Kraken 2. Genome Biology 2019, 20, 257.

(8) Blanco-Míguez, A., et al. Extending and improving metagenomic taxonomic profiling with uncharacterized species using MetaPhlAn 4. Nature Biotechnology 2023, 41, 1633–1644.

(9) Penzlin, A.; Lindner, M. S.; Doellinger, J.; Dabrowski, P. W.; Nitsche, A.; Renard, B. Y. Pipasic: similarity and expression correction for strain-level identification and quantification in metaproteomics. Bioinformatics 2014, 30, i149–i156.

(10) Nesvizhskii, A. I.; Aebersold, R. Interpretation of Shotgun Proteomic Data: The Protein Inference Problem. Molecular & Cellular Proteomics 2005, 4, 1419–1440.

(11) Renard, B. Y.; Xu, B.; Kirchner, M.; Zickmann, F.; Winter, D.; Korten, S.; Brattig, N. W.; Tzur, A.; Hamprecht, F. A.; Steen, H. Overcoming Species Boundaries in Peptide Identification with Bayesian Information Criterion-driven Error-tolerant Peptide Search (BICEPS) *. Molecular & Cellular Proteomics 2012, 11, M111.014167–12.

(12) Muth, T.; Kohrs, F.; Heyer, R.; Benndorf, D.; Rapp, E.; Reichl, U.; Martens, L.; Renard, B. Y. MPA Portable: A Stand-Alone Software Package for Analyzing Metaproteome Samples on the Go. Analytical Chemistry 2018, 90, 685–689.

(13) Gurdeep Singh, R.; Tanca, A.; Palomba, A.; Van der Jeugt, F.; Verschaffelt, P.; Uz-zau, S.; Martens, L.; Dawyndt, P.; Mesuere, B. Unipept 4.0: functional analysis of metaproteome data. Journal of Proteome Research 2018,

(14) Cheng, K.; Ning, Z.; Zhang, X.; Li, L.; Liao, B.; Mayne, J.; Figeys, D. MetaLab 2.0 Enables Accurate Post-Translational Modifications Profiling in Metaproteomics. Journal of the American Society for Mass Spectrometry 2020, 31, 1473–1482.

(15) Riffle, M.; May, D.; Timmins-Schiffman, E.; Mikan, M.; Jaschob, D.; Noble, W.; Nunn, B.; Riffle, M.; May, D. H.; Timmins-Schiffman, E.; Mikan, M. P.; Jaschob, D.; Noble, W. S.; Nunn, B. L. MetaGOmics: A Web-Based Tool for Peptide-Centric Functional and Taxonomic Analysis of Metaproteomics Data. Proteomes 2017, 6, 2.

(16) Schiebenhoefer, H.; Schallert, K.; Renard, B. Y.; Trappe, K.; Schmid, E.; Benndorf, D.; Riedel, K.; Muth, T.; Fuchs, S. A complete and flexible workflow for metaproteomics data analysis based on MetaProteomeAnalyzer and Prophane. Nature Protocols 2020,

(17) Buchfink, B.; Reuter, K.; Drost, H.-G. Sensitive protein alignments at tree-of-life scale using DIAMOND. Nature Methods 2021, 18, 366–368, Bandiera abtest: a Cc license type: cc by Cg type: Nature Research Journals Number: 4 Primary atype: Research Subject term: Computational biology and bioinformatics;Genome informatics;Genomic analysis;Sequencing;Software Subject term id: computational-biology-and-bioinformatics;genome-informatics;genomic-analysis;sequencing;software.

(18) Steinegger, M.; Söding, J. MMseqs2 enables sensitive protein sequence searching for the analysis of massive data sets. Nature Biotechnology 2017, 35, 1026–1028, 00026.

(19) Lin, H.; Peddada, S. D. Analysis of compositions of microbiomes with bias correction. Nature Communications 2020, 11, 3514.

(20) Fernandes, A. D.; Reid, J. N.; Macklaim, J. M.; McMurrough, T. A.; Edgell, D. R.; Gloor, G. B. Unifying the analysis of high-throughput sequencing datasets: characterizing RNA-seq, 16S rRNA gene sequencing and selective growth experiments by compositional data analysis. Microbiome 2014, 2, 15.

(21) Mallick, H. et al. Multivariable association discovery in population-scale meta-omics studies. PLOS Computational Biology 2021, 17, e1009442.

(22) Piro, V. C.; Lindner, M. S.; Renard, B. Y. DUDes: a top-down taxonomic profiler for metagenomics. Bioinformatics 2016, 32, 2272–2280.

(23) The UniProt Consortium UniProt: a worldwide hub of protein knowledge. Nucleic Acids Research 2019, 47, D506–D515.

(24) Lloyd-Price, J.; Mahurkar, A.; Rahnavard, G.; Crabtree, J.; Orvis, J.; Hall, A. B.; Brady, A.; Creasy, H. H.; McCracken, C.; Giglio, M. G.; McDonald, D.; Franzosa, E. A.; Knight, R.; White, O.; Huttenhower, C. Strains, functions and dynamics in the expanded Human Microbiome Project. Nature 2017, 550, 61–66.

(25) Blaxter, M.; Pauperio, J.; Schoch, C.; Howe, K. Taxonomy Identifiers (TaxId) for Biodiversity Genomics: a guide to getting TaxId for submission of data to public databases. Wellcome Open Research 2024, 9, 591.

(26) Nesvizhskii, A. I.; Keller, A.; Kolker, E.; Aebersold, R. A Statistical Model for Identifying Proteins by Tandem Mass Spectrometry. Analytical Chemistry 2003, 75, 4646–4658.

(27) Schulze, S.; Igiraneza, A. B.; Kösters, M.; Leufken, J.; Leidel, S. A.; Garcia, B. A.; Fufezan, C.; Pohlschroder, M. Enhancing Open Modification Searches via a Combined Approach Facilitated by Ursgal. Journal of Proteome Research 2021, 20, 1986–1996.

(28) Kleiner, M.; Thorson, E.; Sharp, C. E.; Dong, X.; Liu, D.; Li, C.; Strous, M. Assessing species biomass contributions in microbial communities via metaproteomics. Nature Communications 2017, 8, 1558, 00007.

(29) Perez-Riverol, Y.; Bandla, C.; Kundu, D.; Kamatchinathan, S.; Bai, J.; Hewapathirana, S.; John, N.; Prakash, A.; Walzer, M.; Wang, S.; Vizcáıno, J. The PRIDE database at 20 years: 2025 update. Nucleic Acids Research 2025, 53, D543–D553.

(30) Kong, A. T.; Leprevost, F. V.; Avtonomov, D. M.; Mellacheruvu, D.; Nesvizhskii, A. I. MSFragger: ultrafast and comprehensive peptide identification in mass spectrome-try–based proteomics. Nature Methods 2017, 14, 513–520.

(31) Yu, F.; Haynes, S. E.; Nesvizhskii, A. I. IonQuant Enables Accurate and Sensitive Label-Free Quantification With FDR-Controlled Match-Between-Runs. Molecular & Cellular Proteomics 2021, 20.

(32) da Veiga Leprevost, F.; Haynes, S. E.; Avtonomov, D. M.; Chang, H.-Y.; Shanmugam, A. K.; Mellacheruvu, D.; Kong, A. T.; Nesvizhskii, A. I. Philosopher: a versatile toolkit for shotgun proteomics data analysis. Nature Methods 2020, 17, 869–870.

(33) Verschaffelt, P.; Van Thienen, P.; Van Den Bossche, T.; Van der Jeugt, F.; De Tender, C.; Martens, L.; Dawyndt, P.; Mesuere, B. Unipept CLI 2.0: adding support for visualizations and functional annotations. Bioinformatics 2020, 36, 4220–4221.

(34) Grossegesse, M.; Horn, F.; Kurth, A.; Lasch, P.; Nitsche, A.; Doellinger, J. vPro-MS enables identification of human-pathogenic viruses from patient samples by untargeted proteomics. Nature Communications 2025, 16, 7041.

